# Detection of *Mycobacterium leprae* DNA in soil; Multiple needles in the haystack?

**DOI:** 10.1101/367219

**Authors:** Maria Tió-Coma, Thomas Wijnands, Louise Pierneef, Anna Katarina Schilling, Korshed Alam, Johan Chandra Roy, William R. Faber, Henk Menke, Toine Pieters, Karen Stevenson, Jan Hendrik Richardus, Annemieke Geluk

## Abstract

**Background:** Leprosy is an infectious disease caused by *Mycobacterium leprae* and *Mycobacterium lepromatosis* affecting the skin and nerves. Despite decades of availability of adequate treatment, transmission is unabated and routes of transmission are not completely understood. Notwithstanding the general assumption that untreated *M. leprae* infected humans represent the major source of transmission, scarce reports indicate that environmental specimens could play a role as a reservoir as well.

**Methodology:** In order to identify patterns of bacterial transmission, this study investigated whether *M. leprae* DNA is present in soil of regions where leprosy is endemic or areas with possible animal reservoirs (armadillos and red squirrels). Soil samples were collected in Bangladesh, Suriname and the British Isles. The presence of *M. leprae* DNA was determined by RLEP PCR and *M. leprae* SNP types were further identified by Sanger sequencing of loci 1-3.

**Results:** *M. leprae* DNA was identified in soil samples from environments inhabited by leprosy patients (Bangladesh), armadillos (Suriname) and the lepromatous Eurasian red squirrels (British Isles). In Bangladesh SNP type 1 was detected, Suriname soil contained SNP type 1 or 2, whilst SNP typing was not feasible for the British samples.

**Conclusions:** It is conceivable that, besides humans and animals, environmental reservoirs may play a role in transmission. Frequent, intense contact with multibacillary leprosy patients poses the highest risk of transmission, and even though the risk of environmental contamination is low, it may offer a possible explanation for the occurrence of leprosy in individuals in areas without any reported human leprosy.

## Introduction

Leprosy is a debilitating infectious disease caused by *Mycobacterium leprae* and *Mycobacterium lepromatosis* that is still considered a major threat in developing countries by WHO, remaining persistently endemic in regions in Africa, South America and Asia. Every year more than 200,000 new patients are still diagnosed and this new case detection rate has been virtually stable over the last decade(1). These facts indicate that multidrug therapy (MDT), although effective to treat leprosy, is insufficient to prevent transmission(2).

Granting *M. leprae* transmission is not completely understood, risk factors for development of leprosy have been identified including close contact with untreated, multibacillary patients(3), human susceptibility genes(4, 5), infection with soil transmitted helminths(6), as well as food shortage(7).

No studies exist that unequivocally demonstrate the mechanisms by which bacteria travel from one organism to another. However, based on existing evidence, skin-to-skin contact, aerosols as well as shedding of bacteria into the environment subsequently followed by infection of other individuals remain the most obvious options for human leprosy(8), (9). Still these routes provide no explanation for the occurrence of leprosy in individuals without known contact to leprosy patients or in areas without any reported new cases(8, 10).

Through PCR amplification of *M. leprae* DNA, its presence has been detected in environmental samples such as soil(11, 12) and water(13) in areas inhabited by leprosy patients in Brazil and India. The viability of *M. leprae* was assessed by its multiplication in footpads of wild type mice and showed that *M. leprae* can remain alive in wet soil for 46 days(14). Moreover, viability of *M. leprae* bacilli in soil from India has been studied by 16S ribosomal RNA gene analysis(15). This study showed that 25% of the soil samples collected from patients’ areas contained *M. leprae* 16S ribosomal RNA, suggesting the presence of viable *M. leprae* in the soil. Additionally, *M. leprae* can survive within environment–free living amoebic cysts up to 8 months(16).

Recently, *M. leprae* and *M. lepromatosis* were identified in red squirrels from the British Isles causing lepromatous disease in several animals^(17, 18)^. Phylogenetic analyses determined that the *M. leprae* strain in squirrels (3I) was related to the lineage circulating in Medieval England, suggesting the red squirrels as a contemporary reservoir of the bacilli.

Zoonotic transmission of *M. leprae* from armadillos has been detected in the southeastern United States where wild armadillos and patients were infected with the same genotype (3I-2-v1)^(19)^.

Furthermore, although the prevalence of leprosy in nonhuman primates (NHP) seems to be quite low, *M. leprae* infections have also been reported in NHP^(20)^ carrying *M. leprae* strains closely related to the human strains, suggesting that NHPs transmission can occur from human (or human sources like trash), but also among NHPs(20).

In this study, we aimed to explore whether soil could be a potential environmental reservoir of *M. leprae*. For this purpose, we investigated the presence of *M. leprae* DNA in soil from regions with varying human leprosy endemicity in Bangladesh, Suriname, Brownsea Island and the Isle of Arran (17).

## Materials and methods

### DNA extraction from soil

Moist soil samples from 3 regions (Table 1) were collected at a depth of 2–8 cm in areas without sun light and stored in 50 ml tubes (Greiner Bio-One, Kremsmünster, Austria): in Bangladesh close to the bedroom of leprosy patients’ homes (n=25) and from areas without known leprosy patients (n=2); in Suriname (Batavia and Groot Chatillon (former leprosy colonies), Pikin Slee and Gujaba) from areas known to be inhabited by nine-banded armadillos (n=28) (samples Suriname 2, 3 and 6 from Batavia and Groot Chatillon were previously described (van Dissel et al. submitted) and are presented here for reference purposes); in the British Isles in the habitat of Eurasian red squirrels carrying *M. leprae* (Brownsea Island, n=10) and *M. lepromatosis* (Isle of Arran, n=10). As a negative control soil was obtained from the surroundings of the Leiden University Medical Centre (The Netherlands). As a positive control, the negative control soil was spiked with 10^8^ cells of *M. leprae* NHPD-63.

**Table 1.** Origin, number and location of soil samples.

| Origin | Number | Area of collection |
| --- | --- | --- |
| Bangladesh | 25 | Houses of leprosy patients |
| Bangladesh | 2 | Area without any reported case of leprosy |
| Suriname | 28 | Surroundings of armadillos' habitats |
| British Isles | 20 | Areas frequented by red squirrels infected with *M. leprae* (Brownsea Island) or *M. lepromatosis* (Arran Isle) |
| The Netherlands | 2 | Control soil |

Summary of soil collected and brief description of area.

DNA was extracted from 10 g of soil using DNeasy PowerMax Soil (Qiagen, Valencia, CA) as per manufacturer’s instructions.

### PCR amplification of RLEP and LPM244

To detect the presence of *M. leprae* DNA in soil, a PCR amplifying an *M*. leprae-specific repetitive sequence (RLEP) was performed. PCR amplification of a 129 bp sequence of RLEP(21) was carried by addition of 10 μl 5x Gotaq^®^ Flexi buffer (Promega, Madison, WI), 5 μl MgCl_2_ (25 mM), 2 μl dNTP mix (5 mM), 0.25 μl Gotaq^®^ G2 Flexi DNA Polymerase (5 u/μl), 5 μl (2 μM) forward and reverse primers (Supplementary Table 1) and 5 μl template DNA in a final volume of 50 μl. DNA from *M. bovis* BCG P3 and *M. tuberculosis* H37Rv were used to assess PCR-specificity. As PCR positive controls DNA from *M. leprae* Br4923 and Thai-53 were used.

To detect inhibition of PCR due to remaining soil components, 1 μl of *M. leprae* DNA was added to the aforementioned PCR mixes together with 5 μl template DNA. In samples presenting PCR inhibition, 5 μl (2mM) Bovine Serum Albumin (BSA) Fraction V (Roche Diagnostics, Indianapolis, IN) were added to the PCR mixes.

PCR mixes were denatured for 2 min at 95 °C followed by 40 cycles of 30 s at 95 °C, 30 s at 65°C and 30 s at 72 °C and a final extension of 10 min at 72 °C. PCR products (15μl) were used for electrophoresis in a 3.5% agarose gel at 130V. Amplified DNA was visualized by Midori Green Advance staining (Nippon Genetics Europe, Dueren, Germany) using a Gel Doc System (Bio-Rad Laboratories, Hercules, CA).

PCR to detect *M. lepromatosis* was performed for soil from the British Isles. The primers (LPM244) amplify a 244 bp region of the *hemN* gene not present in *M. leprae* or other mycobacteria(22). PCR was performed as explained above with LMP244 primers (Supplementary Table 1) and an annealing temperature of 53 °C. *M. lepromatosis* DNA was used as a positive control.

### SNP typing

To determine the SNP type (1, 2, 3 or 4) of *M. leprae*, SNP-14676 (locus 1), SNP-1642875 (locus 2) and SNP-2935685 (locus 3) were amplified and sequenced as described(23) with minor modifications: PCRs were performed with 5 μl of template DNA using the aforementioned PCR mixes and forward and reverse primers for loci 1-3 (Supplementary Table 1) in a final volume of 50 μl. DNA was denatured for 2 minutes at 95°C, following 45 cycles of 30 s at 95°C, 30 s at 58 °C and 30 s at 72 °C and a final extension cycle of 10 min at 72°C. PCR products were resolved by agarose gel electrophoresis as explained above. PCR products showing a band were purified prior to sequencing using the Wizard SV Gel and PCR Clean-Up System (Promega, Madison, WI). Sequencing was performed on the ABI3730xl system (Applied Biosystems, Foster City, CA) using the BigDye Terminator Cycle Sequencing Kit (Thermo Fisher Scientific, Waltham, MA).

## Results

### Detection of *M. leprae* DNA in soil

To determine whether *M. leprae* DNA is present in the environment surrounding the houses of leprosy patients, the habitat of armadillos and red squirrels with leprosy-like disease, soil was collected in each area. PCR amplification of a 129 bp sequence of the RLEP region from *M. leprae* was performed in a total of 75 soil samples from 3 different regions (Table 1). Control soil samples did not show amplification of the fragment in RLEP PCR, whereas the same sample spiked with *M. leprae* bacilli presented a clear band confirming the applicability of the method to isolate, purify and detect *M. leprae* in soil. PCR amplification of 5 μl of *M. bovis* BCG P3 and *M. tuberculosis* H37Rv DNA did not show amplification of RLEP showing specificity of the PCR for *M. leprae* DNA.

In Bangladesh, 4 out of 25 collected samples were positive for RLEP PCR (Fig 1, Table 2; Supplementary Table 2), all of which were collected in houses of leprosy patients with high bacillary load (BI) (Fig 2). *M. leprae* DNA was not detected in the two soil samples from areas in Bangladesh without any reported leprosy cases (Supplementary Fig 1).

**Fig 1.**
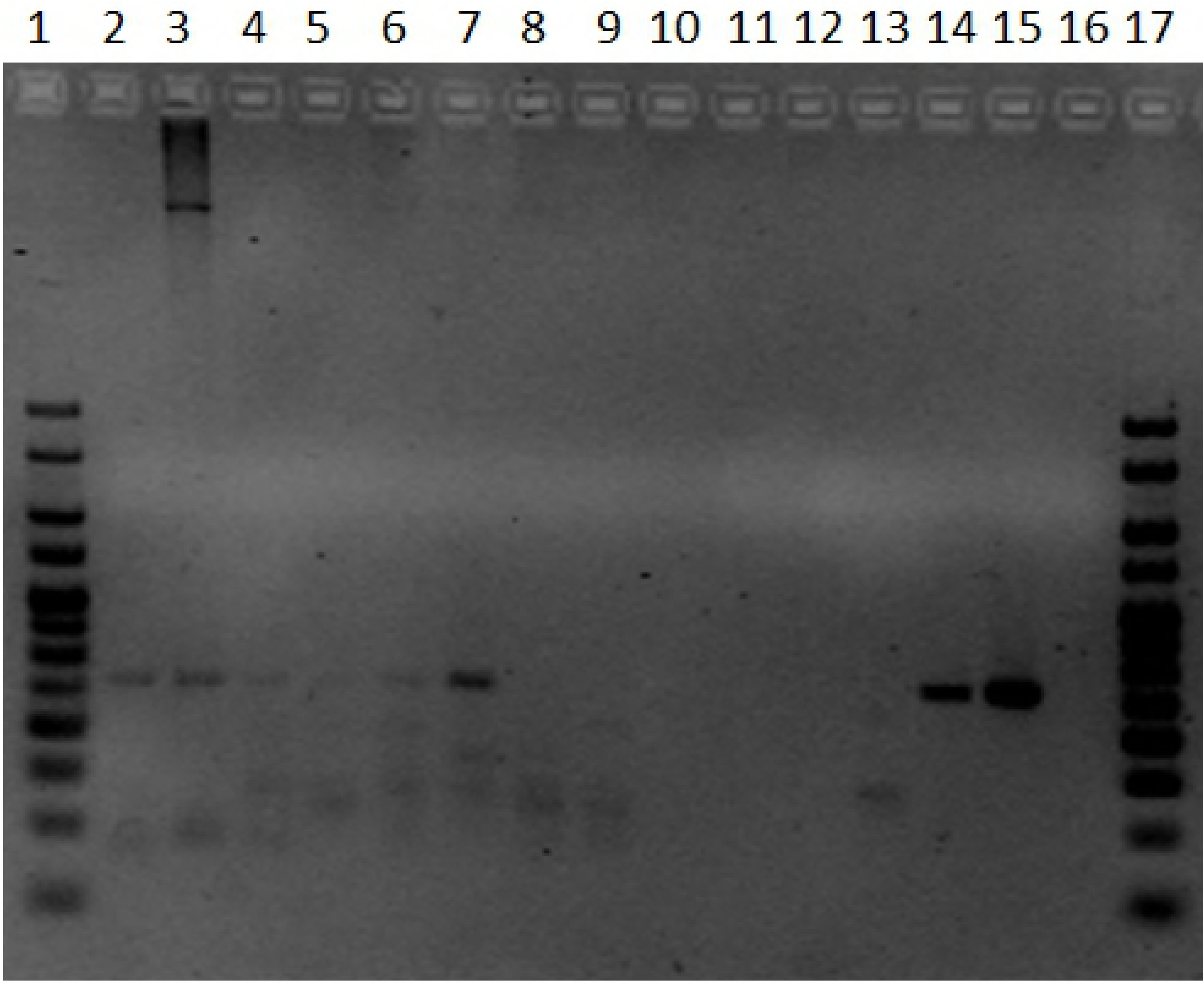
Gel of PCR for RLEP region to detect presence of M. leprae in soil samples. PCR products were electrophoresed in a 3.5% agarose gel. The size of the amplified RLEP sequence is 129 bp. Lanes 2 to 4 represent soil samples collected in Suriname (Suriname 2, 3, and 6), lanes 5 to 14 are soil samples collected in Bangladesh (01/65959/00, 01/65922/00, 01/65958/00, 02/65971/00, 02/22705/00, 01/65945/00, 01/65942/00, 01/65975/00, 01/22711/00 and 01/22723/00), lane 15 is DNA of *M. leprae* Thai-53 strain, lane 16 is a negative PCR control and lanes 1 and 17 are 25 bp HyperLadder (Bioline, Taunton, MA).

**Fig 2.**
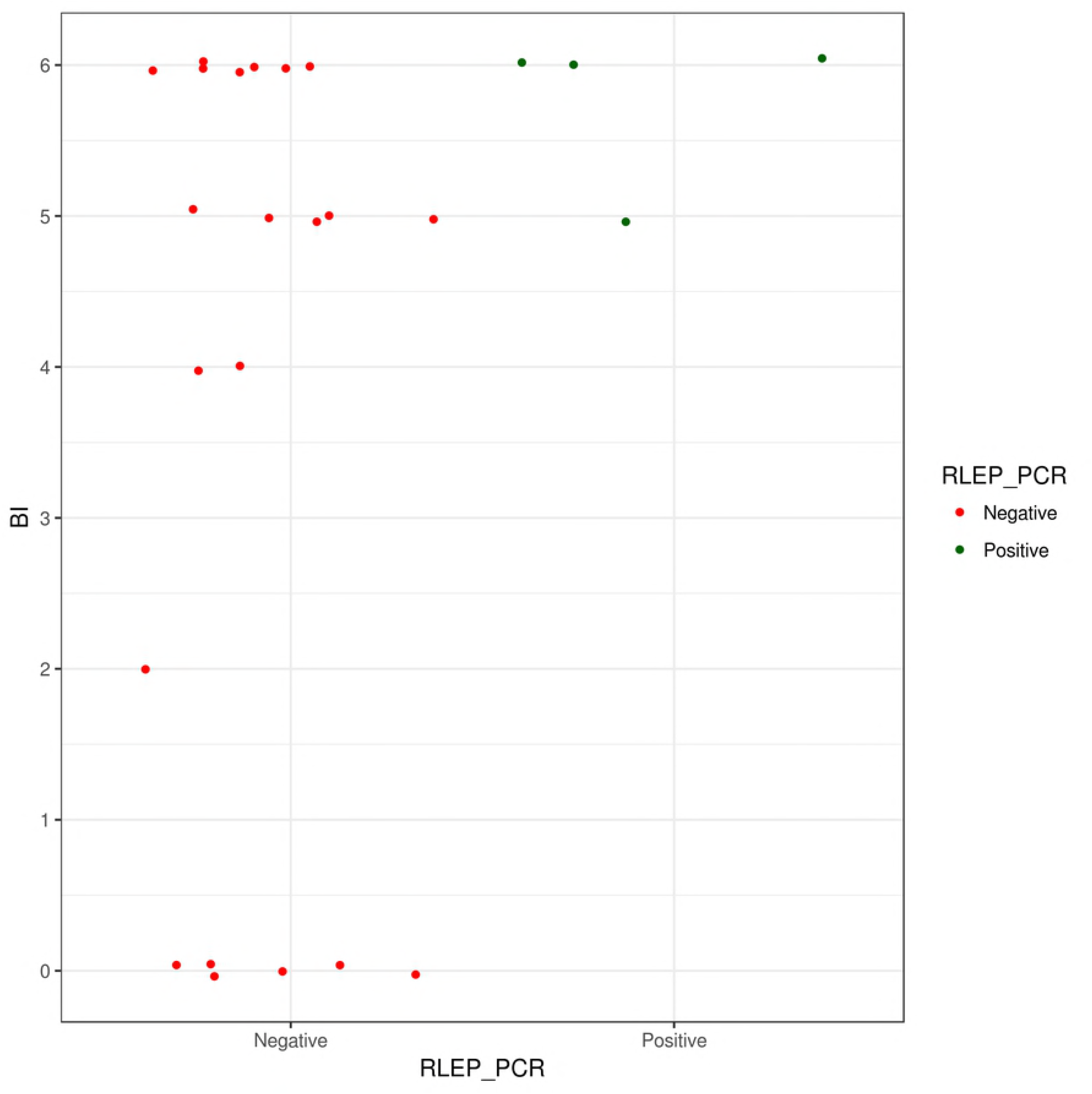
RLEP PCR positivity in soil samples from Bangladesh and bacillary load (BI) of patient. Soil samples collected in Bangladesh are represented in the graph by dots and sorted based on RLEP PCR results and bacillary load of the patient living in the household where the soil was collected. In Suriname, samples (n=28) were taken in three different locations inhabited by armadillos and *M. leprae* DNA was detected in 3 samples obtained at former leprosy colonies in Batavia and Groot-Chatillon (Fig 1, Table 2; Supplementary Table 3).

**Table 2.** RLEP PCR results for M. leprae DNA derived from soil samples. RLEP PCR result to detect *M. leprae* DNA in soil samples from Bangladesh, Suriname, Brownsea Island and Isle of Arran. A positive result is determined by a visible band of 129 bp in an agarose gel.

| Origin | Positive |  | Negative |  |
| --- | --- | --- | --- | --- |
|  | Number | % | Number | % |
| Bangladesh | 4 | 16.0 | 21 | 84.0 |
| Suriname | 3 | 10.7 | 25 | 89.3 |
| Brownsea Island | 1 | 10.0 | 9 | 90.0 |
| Isle of Arran | 0 | 0.0 | 10 | 100.0 |

Since all PCRs performed with UK samples were negative, we investigated whether PCRs were inhibited by compounds in the soil. DNA of *M. leprae* was added to the PCR mixes containing the DNA isolated from all soil samples and inhibition of PCR was determined by a negative PCR result. Inhibition was observed in 7 of the 10 soil samples from Brownsea Island, 8 out of the 10 from the Isle of Arran and 1 out of the 28 from Suriname. Since humic acid in soil can act as a PCR inhibitor(24, 25), 5 μl of 2 mM BSA was added to the PCRs with soil samples from the British Isles to overcome inhibition. Indeed, addition of BSA to soil-DNA spiked with *M. leprae* DNA (Br4923 or Thai-53), resulted in PCR-positivity for all spiked samples, indicating that BSA can prevent PCR inhibition due to undetermined soil compounds (data not shown).

Ten soil samples were collected in the surroundings of the infected red squirrels one of which was RLEP PCR positive (Tables 2 and 3). To determine whether *M. lepromatosis* DNA was also present in soil from the Isle of Arran with reported *M. lepromatosis* infection in red squirrels, PCRs were performed amplifying a 244 bp region of the *hemN* gene unique of *M. lepromatosis* (22). None of the 10 soil samples collected resulted in PCR-positivity using LPM244 primers.

**Table 3.** *SNP type results*. Polymorphic sites in the genome of *M. leprae:* locus 1 (SNP-14676), locus 2 (SNP-1642875) and locus 3 (SNP-2935685) and the corresponding SNP type. Nucleic acid corresponding to each polymorphic site of *M. leprae* reference strains Tamil Nadu and Br4923 and soil samples that were successfully sequenced. When PCR amplification or sequencing of the locus was not successful it is marked as undetermined (UD).

|  | Locus 1 | Locus 2 | Locus 3 | SNP type |
| --- | --- | --- | --- | --- |
| Tamil Nadu (reference strain) | C | G | A | 1 |
| Br4923 (reference strain) | T | T | C | 4 |
| Suriname 2 | UD | UD | A | 1 or 2 |
| Suriname 3 | UD | UD | A | 1 or 2 |
| Suriname 6 | C | UD | A | 1 or 2 |
| Bangladesh 01/65922/00 | UD | G | UD | 1 |
| Bangladesh 01/65958/00 | UD | G | UD | 1 |
| Bangladesh 01/22723/00 | C | G | A | 1 |

Next, for all RLEP PCR positive samples from Bangladesh (n=4), Suriname (n=3) and the British Isles (n=1) the PCR-amplified 129 bp RLEP region was sequenced. Sequence alignment with the RLEP region of *M. leprae* was found for all 8 samples, confirming that *M. leprae* specific DNA can be identified in soil using the above described procedure.

### SNP typing

SNP types of the 8 RLEP PCR positive soil samples were investigated and determined according to the combination of SNPs in loci 1-3 as described by Monot *et al*.(23) RLEP-positive soil from Bangladesh were typed as SNP type 1 (Table 3) according to the polymorphism in locus 2 or loci 1-3 (01/22723/00, Fig 3). For the soil from Suriname the SNP type was narrowed down to either SNP type 1 or 2 since only sequencing of locus 3 (Suriname 2, 3 and 6) and locus 1 (Suriname 6) were identified. For the RLEP positive sample from Brownsea Island it was not possible to obtain sequence information for any of the polymorphic loci to assign a SNP type. This was most likely due to the small amount of *M.leprae* DNA in the samples.

**Fig 3.**
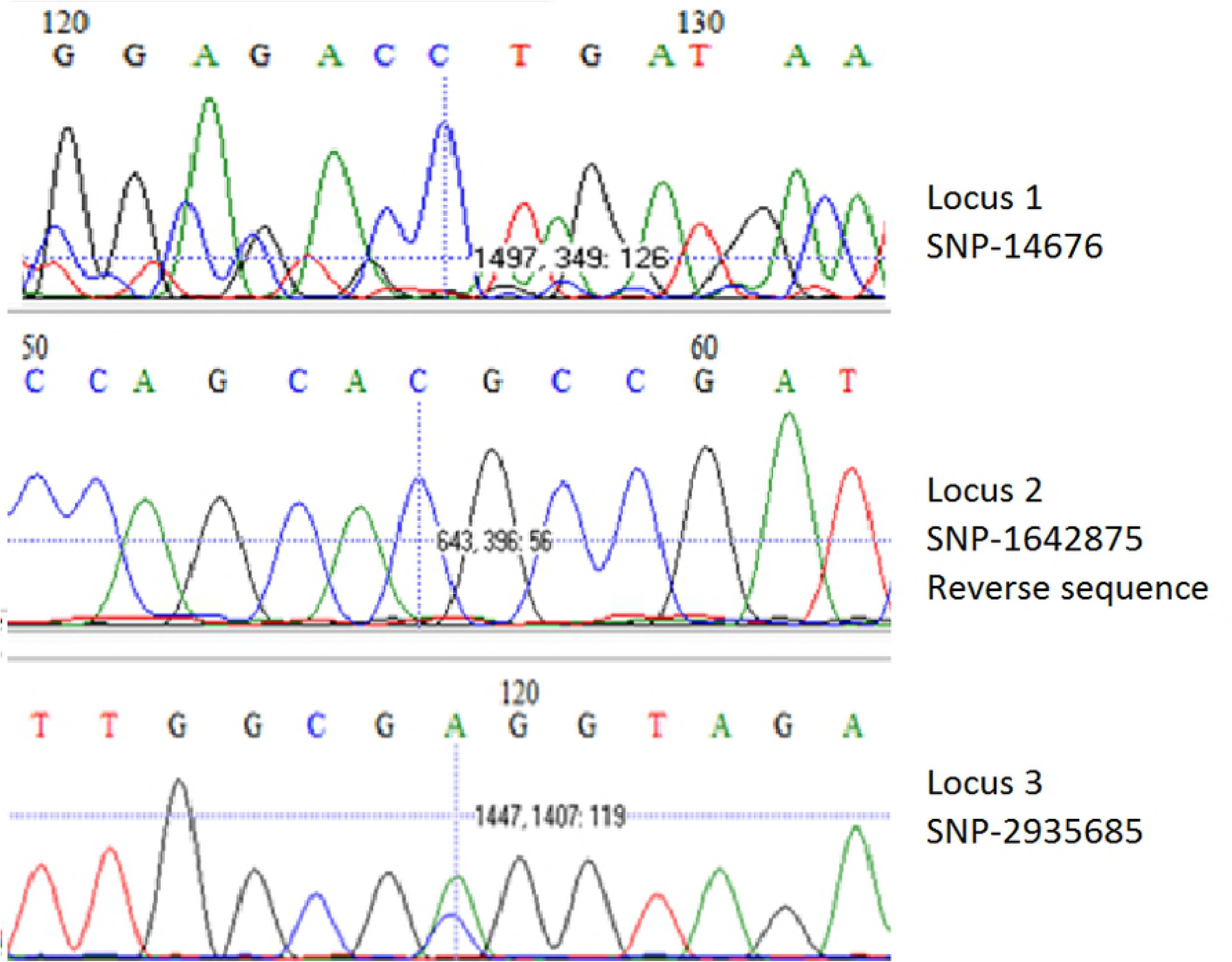
SNP analysis of loci 1, 2 and 3 from a representative *M. leprae* positive soil sample collected in Bangladesh. Sequencing results of locus 1 (SNP-146763) top, reverse sequence of locus 2 (SNP-1642875) middle and locus 3 (SNP-2935685) bottom, from soil sample Bangladesh 01/22723/00 used to determine the SNP type of the *M. leprae* strain identified (SNP type 1). SNP positions are based on the *M. leprae* TN strain. Vertical bars indicate the polymorphic base.

## Discussion

Human leprosy still poses a considerable health threat in developing countries where transmission is generally assumed to take place via aerosol droplets from nasal cavities of untreated *M. leprae* infected individuals to their close contacts(8, 9). However, nonhuman animal and environmental sources have also been suggested to play a role in the pathogen’s dissemination(8). As paleopathological evidence of leprosy in pre-Columbian America is lacking, leprosy was very likely introduced to the continent by European colonists or the African slave route(23) also resulting in transmission to armadillos. However, nowadays infected armadillos may even be responsible for new cases in human individuals who have never had contact with leprosy patients nor travelled to leprosy endemic areas(10, 26).

In this study, *M. leprae* DNA was identified in soil surrounding leprosy patients’ houses and the habitats of armadillos and red squirrels. However, this study did not asses viability of the bacteria. Hence, soil may represent a (temporary) reservoir for *M. leprae* contributing to transmission and infection of humans and animals.

Understanding how *M. leprae* is transmitted, and identifying sources of infection is crucial to prevent new cases and thus blocking transmission is essential to ultimately eradicate leprosy.

Although human leprosy was eradicated from the British Isles centuries ago, Eurasian red squirrels have remained a reservoir for *M. leprae*, containing a strain closely related to the strain present in Medieval England (3I). This indicates that *M. leprae* may have persisted in the environment after the human reservoir disappeared. However, *M. leprae* DNA was not abundantly present in soil, suggesting that the risk of environmental contamination is low.

Detection of *M. lepromatosis* DNA by LPM244 PCR is less sensitive than *M. leprae* DNA by RLEP PCR because the genome of *M. lepromatosis* contains only one copy of the *hemN* gene detected by LPM244 compared to 37 copies(27) of the RLEP region(28) in *M. leprae*. Added to the fact that *M. lepromatosis* prevalence in the squirrel population is low, it is therefore possible that sensitivity was not sufficient to detect *M. lepromatosis*.

In Bangladesh, *M. leprae* was only found in soil collected in the houses of patients with high BI index (Fig 2). At those locations more bacteria are shed and thus the likelihood of encountering bacteria in the soil is higher. However, a high BI index of the patient where the soil sample was collected was not necessarily associated with a positive RLEP PCR result. The higher percentage of RLEP positive soil in Bangladesh is likely due to a more targeted selection of the sample location in the houses of leprosy patients as well as the higher leprosy prevalence.

In previous phylogeographic analysis SNP type 1 was identified as the predominant strain type in South Asia(29, 30) and was likely introduced to South Asia from other parts of that continent(30). The SNP type found in soil samples from Bangladesh (SNP type 1) is therefore in accordance with previous phylogeographic data(29).

In summary, this study demonstrates the presence of *M. leprae* DNA in soil, contributing to a OneHealth view on transmission including humans, animals and the environment. Further research is needed, however, to confirm whether *M. leprae* DNA in soil is derived from viable bacteria that can survive in smaller hosts such as helminths or amoebas. Thus, strategies aimed at prevention of transmission by administration of post-exposure prophylaxis to infected individuals should, besides human reservoirs of *M. leprae*, also consider environmental sources of (re)infection.

## Acknowledgements

The authors gratefully acknowledge all patients and control participants. LUMC and TLMI,B are part of the IDEAL (***I***nitiative for ***D***iagnostic and ***E***pidemiological ***A***ssays for ***L***eprosy) Consortium.

We thank Dr. L. Adams (Louisiana State University, LA) for providing *M. leprae* cells and stimulating discussions and Prof. Xiang-Yang Han (MD Anderson Cancer Center, TX, USA) for providing *M. lepromatosis* DNA.

## References

1. World Health Organization. Global leprosy update, 2016: accelerating reduction of disease burden. 92: WHO; 2017. p. 501–20.

2. Blok DJ, de Vlas SJ, Geluk A, Richardus JH. Minimum requirements and optimal testing strategies of a diagnostic test for leprosy as a tool towards zero transmission: A modeling study. PLOS Neglected Tropical Diseases. 2018;12(5):e0006529.

3. Moet FJ, Meima A, Oskam L, Richardus JH. Risk factors for the development of clinical leprosy among contacts, and their relevance for targeted interventions. Leprosy review. 2004;75(4):310–26.

4. Liu H, Irwanto A, Fu X, Yu G, Yu Y, Sun Y, et al. Discovery of six new susceptibility loci and analysis of pleiotropic effects in leprosy. Nature genetics. 2015;47(3):267–71.

5. Alter A, Grant A, Abel L, Alcais A, Schurr E. Leprosy as a genetic disease. Mammalian genome: official journal of the International Mammalian Genome Society. 2011;22(1–2): 19–31.

6. Hagge DA, Parajuli P, Kunwar CB, Rana D, Thapa R, Neupane KD, et al. Opening a Can of Worms: Leprosy Reactions and Complicit Soil-Transmitted Helminths. EBioMedicine. 2017;23:119–24.

7. Feenstra SG, Nahar Q, Pahan D, Oskam L, Richardus JH. Recent food shortage is associated with leprosy disease in Bangladesh: a case-control study. PLOS Neglected Tropical Diseases. 2011;5(5):e1029.

8. Araujo S, Freitas LO, Goulart LR, Goulart IM. Molecular evidence for the aerial route of infection of *Mycobacterium leprae* and the role of asymptomatic carriers in the persistence of leprosy. Clinical infectious diseases: an official publication of the Infectious Diseases Society of America. 2016;63(11):1412–20.

9. Bratschi MW, Steinmann P, Wickenden A, Gillis TP. Current knowledge on Mycobacterium leprae transmission: a systematic literature review. Leprosy review. 2015;86(2):142–55.

10. Bonnar PE, Cunningham NP, Boggild AK, Walsh NM, Sharma R, Davis IRC. Leprosy in Nonimmigrant Canadian Man without Travel outside North America, 2014. Emerging infectious diseases. 2018;24(1):165–6.

11. Lavania M, Katoch K, Sachan P, Dubey A, Kapoor S, Kashyap M, et al. Detection of Mycobacterium leprae DNA from soil samples by PCR targeting RLEP sequences. The Journal of communicable diseases. 2006;38(3):269–73.

12. Turankar RP, Lavania M, Chaitanya VS, Sengupta U, Darlong J, Darlong F, et al. Single nucleotide polymorphism-based molecular typing of *M. leprae* from multicase families of leprosy patients and their surroundings to understand the transmission of leprosy. Clinical microbiology and infection: the official publication of the European Society of Clinical Microbiology and Infectious Diseases. 2014;20(3):O142–9.

13. Holanda MV, Marques LEC, Macedo MLB, Pontes MAA, Sabadia JAB, Kerr L, et al. Presence of *Mycobacterium leprae* genotype 4 in environmental waters in Northeast Brazil. Revista da Sociedade Brasileira de Medicina Tropical. 2017;50(2):216–22.

14. Desikan KV, Sreevatsa. Extended studies on the viability of *Mycobacterium leprae* outside the human body. Leprosy review. 1995;66(4):287–95.

15. Mohanty PS, Naaz F, Katara D, Misba L, Kumar D, Dwivedi DK, et al. Viability of *Mycobacterium leprae* in the environment and its role in leprosy dissemination. Indian journal of dermatology, venereology and leprology. 2016;82(1):23–7.

16. Wheat WH, Casali AL, Thomas V, Spencer JS, Lahiri R, Williams DL, et al. Long-term survival and virulence of *Mycobacterium leprae* in amoebal cysts. PLoS Neglected Tropical Diseases. 2014;8(12):e3405.

17. Avanzi C, Del-Pozo J, Benjak A, Stevenson K, Simpson VR, Busso P, et al. Red squirrels in the British Isles are infected with leprosy bacilli. Science (New York, NY). 2016;354(6313):744–7.

18. Simpson V, Hargreaves J, Butler H, Blackett T, Stevenson K, McLuckie J. Leprosy in red squirrels on the Isle of Wight and Brownsea Island. The Veterinary record. 2015;177(8):206–7.

19. Truman RW, Singh P, Sharma R, Busso P, Rougemont J, Paniz-Mondolfi A, et al. Probable zoonotic leprosy in the Southern United States. The New England journal of medicine. 2011;364(17):1626–33.

20. Honap TP, Pfister LA, Housman G, Mills S, Tarara RP, Suzuki K, et al. *Mycobacterium leprae* genomes from naturally infected nonhuman primates. PLOS Neglected Tropical Diseases. 2018;12(1):e0006190.

21. Donoghue HD, Holton J, Spigelman M. PCR primers that can detect low levels of *Mycobacterium leprae* DNA. Journal of medical microbiology. 2001;50(2):177–82.

22. Vera-Cabrera L, Escalante-Fuentes W, Ocampo-Garza SS, Ocampo-Candiani J, Molina-Torres CA, Avanzi C, et al. *Mycobacterium lepromatosis* infections in Nuevo León, Mexico. Journal of clinical microbiology. 2015;53(6):1945–6.

23. Monot M, Honore N, Garnier T, Araoz R, Coppee JY, Lacroix C, et al. On the origin of leprosy. Science (New York, NY). 2005;308(5724):1040–2.

24. Tsai YL, Olson BH. Rapid method for separation of bacterial DNA from humic substances in sediments for polymerase chain reaction. Applied and Environmental Microbiology. 1992;58(7):2292–5.

25. Watson RJ, Blackwell B. Purification and characterization of a common soil component which inhibits the polymerase chain reaction. Canadian journal of microbiology. 2000;46(7):633–42.

26. da Silva MBP, Juliana M., Li W, Jackson MG-J, Mercedes, Belisle JT, Bouth RC, Gobbo ARB, Josafá G., et al. Evidence of zoonotic leprosy in Pará, Brazilian Amazon, and increased anti-PGL-I titer in individuals who consume armadillos in their diet. PLOS Neglected Tropical Diseases. 2018;12(6):e0006532.

27. Cole ST, Supply P, Honoré N. Repetitive sequences in *Mycobacterium leprae* and their impact on genome plasticity. Leprosy review. 2001;72(4):449–61.

28. Braet S, Vandelannoote K, Meehan CJ, Brum Fontes AN, Hasker E, Rosa PS, et al. The repetitive element RLEP is a highly specific target for detection of *Mycobacterium leprae*. Journal of clinical microbiology. 2018;56(3).

29. Monot M, Honore N, Garnier T, Zidane N, Sherafi D, Paniz-Mondolfi A, et al. Comparative genomic and phylogeographic analysis of *Mycobacterium leprae*. Nature genetics. 2009;41(12):1282–9.

30. Benjak A, Avanzi C, Singh P, Loiseau C, Girma S, Busso P, et al. Phylogenomics and antimicrobial resistance of the leprosy bacillus *Mycobacterium leprae*. Nature communications. 2018;9(1):352.

